# Non-invasive real-time imaging of reactive oxygen species (ROS) using multispectral auto-fluorescence imaging technique: a novel tool for redox biology

**DOI:** 10.1101/2020.02.18.955112

**Authors:** Abbas Habibalahi, Mahdieh Dashtbani Moghari, Jared M. Campbell, Ayad G. Anwer, Saabah B. Mahbub, Martin Gosnell, Sonia Saad, Carol Pollock, Ewa M. Goldys

## Abstract

Detecting reactive oxygen species (ROS) that play a critical role as redox modulators and signalling molecules in biological systems currently requires invasive methods such as ROS - specific indicators for imaging and quantification. We developed a non-invasive, real-time, label-free imaging technique for assessing the level of ROS in live cells and thawed cryopreserved tissues that is compatible with *in-vivo* imaging. The technique is based on autofluorescence multispectral imaging (AFMI) carried out in an adapted fluorescence microscope with an expanded number of spectral channels spanning specific excitation (365 nm-495 nm) and emission (420 nm-700 nm) wavelength ranges. We established a strong quantitative correlation between the spectral information obtained from AFMI and the level of ROS obtained from CellROX staining. The results were obtained in several cell types (HeLa, PANC1 and mesenchymal stem cells) and in live kidney tissue. Additioanly, two spectral regimes were considered: with and without UV excitation (wavelengths > 400 nm); the latter being suitable for UV-sensitive systems such as the eye. Data were analyzed by linear regression combined with an optimization method of swarm intelligence. This allowed the calibration of AFMI signals to the level of ROS with excellent correlation (R= 0.84, p=0.00) in the entire spectral range and very good correlation (R= 0.78, p=0.00) in the limited, UV-free spectral range. We also developed a strong classifier which allowed us to distinguish moderate and high levels of ROS in these two regimes (AUC= 0.91 in the entire spectral range and AUC = 0.78 for UV-free imaging). These results indicate that ROS in cells and tissues can be imaged non-invasively, which opens the way to future clinical applications in conditions where reactive oxygen species are known to contribute to progressive disease such as in ophthalmology, diabetes, kidney disease, cancer and neurodegenerative diseases.

## 1. INTRODUCTION

Accurate reactive oxygen species (ROS) quantification and characterization plays an important role in the investigation of metabolism [1, 2], inflammatory responses [3, 4], the pathogenesis of the disease [5, 6], treatment monitoring [7–9][10] and potentially provides prognostic information [11]. Currently, standard ROS measurement uses various molecular indicators such as dichlorofluorescein, a fluorescent probe that reacts intracellularly with peroxidase, or CellROX, a fluorogenic probe for measuring oxidative stress in live cells and tissue [12–14]. However, due to challenges with administration and potential toxicity molecular ROS probes may not be suitable for in-vivo use [12]. To overcome these limitations an imaging technique which can measure ROS levels directly, without any staining, in clinical and research environments, is highly desirable.

In this study, we employed autofluorescence multispectral imaging (AFMI) [15, 16] to non-invasively characterize ROS levels. The autofluorescence emission in cells or tissues originates from endogenous fluorophores including collagen, elastin, tryptophan and reduced nicotinamide adenine dinucleotide (phosphate) (NAD(P)H), flavins ([16, 17]). The autofluorescence emission of some of these, most notably NAD(P)H, depends on its oxidation state. NAD(P)H is fluorescent wheras NAD+ is not, and the relative balance of these two depends on the level of ROS [18, 19]. This allows the status of cellular, mitochondrial or cytosolic ROS to be non-invasively evaluated and monitored [20].

AFMI is a new technology that uses an adapted fluorescence microscope which provides light excitation in several narrow (+/− 5 nm) wavelength ranges and collects the native fluorescence emission of the sample, also at defined wavelength ranges [15, 16]. Each excitation/emission wavelength band combination forms a specific spectral channel and we used a number of such channels (here 18) to collect detailed spectral image information from cells and tissue. Following AFMI, the slide-mounted cells and tissue were stained using CellROX and imaged again to establish the reference ROS values. These were matched with the corresponding AFMI data in each individual cell or tissue section separately.

Non-invasive evaluation of ROS in real-time was achieved by calibrating the cellular AFMI data against the ROS data from CellROX, on a single-cell level. This calibration was based on selected color features extracted from the single-cell AFMI images and the corresponding CellROX images. The data sets were then used to develop classifiers, with a threshold based on the median of CellROX values, allowing the non-invasive determination of whether the ROS level in cells and tissues was moderate or elevated. We have also developed a UV-free version of the AFMI vs ROS (CellROX) calibration and a related UV-free classifier where the UV channels with excitation wavelengths of less than 400 nm were avoided; this is relevant for UV-sensitive applications such as in reproductive medicine and ophthalmology. Taken together these results represent a novel methodology that allows objective, non-invasively determination of ROS levels related to oxidative metabolism or redox dysregulation in cells and tissues from multispectral images using automated technology.

## 2. Methods

### 2.1. Sample preparation and ROS stimulation

In this study we investigated neoplastic cells (HeLa and PANC1), human bone marrow derived mesenchymal stem cells (MSC) (isolated as described previously [21, 22]) and cryopreserved whole kidney tissue sections obtained from 13 week old Balb/c mice, which were exposed to either air or smoke in utero, as we have previously demonstrated that they express increased levels of ROS[23]. Full details of cell culture and ethics approvals are included in Supplementary Material note 1. To stimulate ROS accumulation, cells were exposed to 30µM menadione (Sigma) in the culture media for 0, 30, 60, 90 and 120 minutes. For imaging, cells were plated onto 35mm ibiTreat coverslip bottomed dishes with 500μm grids (Ibidi).

### 2.2. Autofluorescence multispectral imaging (AFMI)

The (AFMI) system used in this study is based on an adapted IX83 Olympus microscope as previously reported [17, 24, 25]. It provides narrow excitation wavelength ranges (±5nm) from high power LEDs and several filter cubes to produce defined spectral regions, which span the excitation (340 nm-510 nm) and emission (420 nm-650 nm) wavelength ranges. In total, 18 distinct spectral channels (*N*_*Ch*_ =18) are available and the samples are imaged in all these channels. These channels cover the spectrum of several fluorophores notably NAD(P)H and flavins, whose concentrations depend on the cellular redox conditions. The spectral details of the channels are shown in Supplementary Table 1. The system also takes differential contrast microscopy (DIC) images. A CCD camera with a high quantum efficiency (Andor IXON 885 EMCCD, Andor Technology Ltd., UK) was used to acquire the fluorescence images in the spectral channels. The images were captured at 40 × magnification. We imaged in the order of 100 cells from each cell type and about 70 tissue regions from the kidney tissue (~500um^2^). The channel images were prepared for subsequent quantitative analysis. Image preparation is a multistep procedure that minimizes sources of errors such as Poisson’s noise, dead or saturated pixels, background fluorescence and the illumination curvature [26]. These artifacts were treated as previously reported [16, 27].

### 2.3. CellROX and nuclear staining

Following AFMI imaging, cell/tissue samples were stained with CellROX Deep Red (Thermo Fisher). CellROX Deep Red is a cell-permeant dye which stains the cytoplasm. This probe detects superoxide anion and hydroxyl radicals [28]. To obtain the reference ROS values, CellROX was applied to cells/tissue on the slides for 30 minutes as per manufacturer’s instructions, followed by imaging on an Olympus FV3000 confocal microscope for hMSC cell lines and kidney tissue, or a Leica FLUOTAR340 for the cancer cell lines. Nuclear counter-staining was performed with Hoechst 3(3342 (NucBlue, Thermo Fisher).

**Figure 1.**
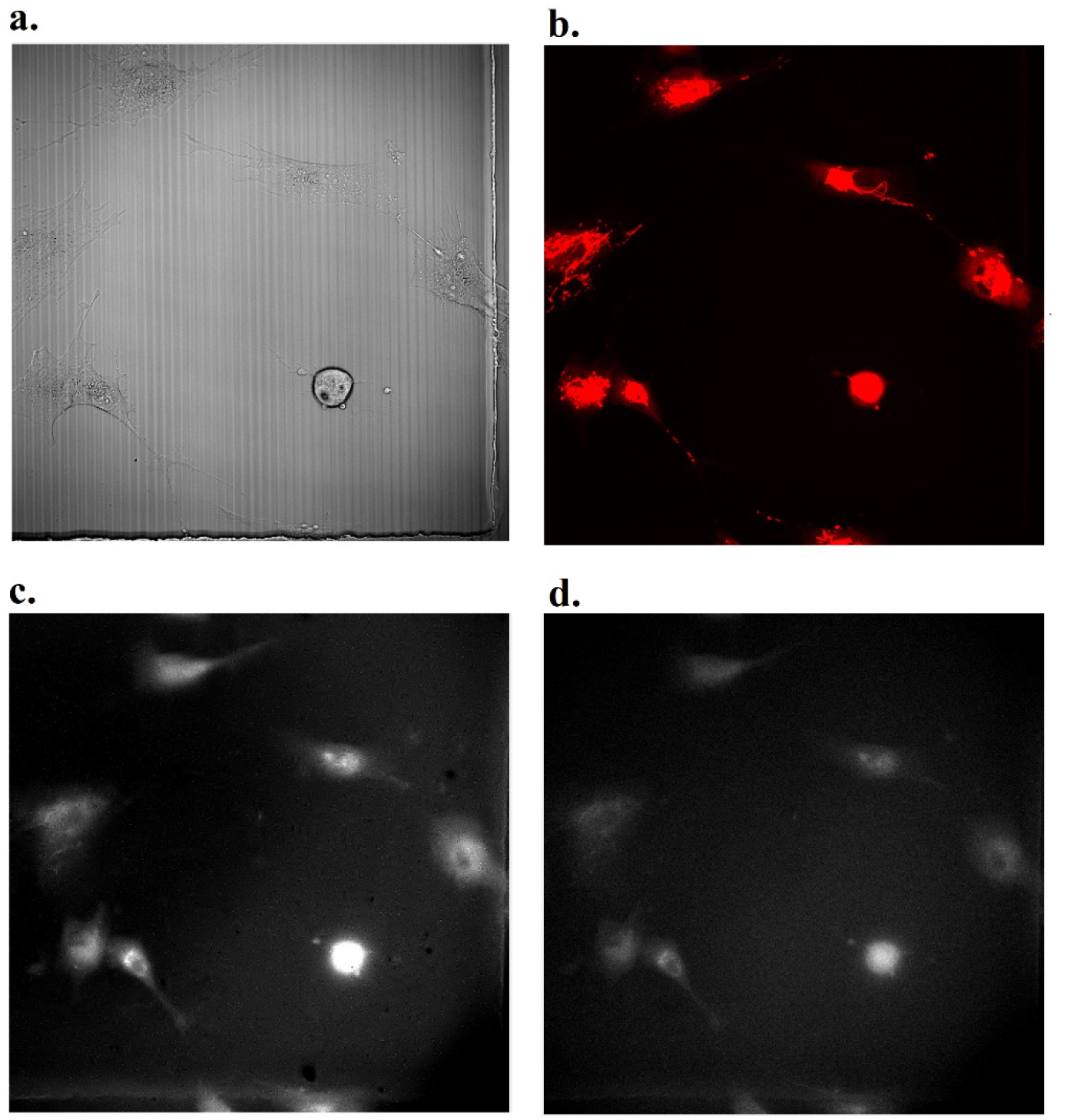
(a) An example of MSC DIC image. (b) an example CellROX image. (c),(d) examples of spectral image channels 2 and 13, respectively.

### 2.4. Image analysis

The cells and tissue regions were segmented manually, on the basis of their DIC images taken during AFMI imaging, and by using confocal CellROX images. Further, the average channel or CellROX values in the segmented cells or tissue regions were calculated. Autofluorescence feature vectors containing these average channel values (18 channels) and all ratio of average channel values (18×17 ratios) for each cell were obtained. The CellROX images were, similarly, segmented and the average cellular value of the CellROX signal (providing the reference ROS value) was found for each cell/tissue sector. Finally, the corresponding cells and tissue regions were identified in the AFMI and CellROX images. The ROS reference values were then appended to the 324-dimensional autofluorescence feature vectors (AFMI vectors) obtained from AFMI for all cells or tissue regions.

### 2.5. Calibration - relating AFMI to ROS reference values from CellROX

The aim of system calibration is to identify a small subset of AFMI features whose optimized linear combination is approximately proportional to the CellROX ROS values, for all individual cells, or tissue sectors. This is achieved by swarm intelligence combined with the regression model. The optimized linear combination of AFMI features (a “spectral variable”) represents the cellular CellROX data in the “best” way, by numerically minimizing the sum of squared errors for all cells. The error is the difference between the average cellular CellROX values and the average values of that spectral variable for a particular cell or tissue region (see Supplementary Material note 2).

### 2.6 Non-invasive assessment of ROS by classification of AFMI data

To distinguish cells with high ROS levels from cells with moderate ROS levels, we developed the AFMI-based classification method. In this method cells and tissue regions were assigned to high or moderate ROS groups using the median cellular value of CellROX signal as a threshold.The cluster analysis was first applied to separate cell groups and further classifiers were employed to predict the pre-defined cell labels [29].

### 2.7 UV-free ROS calibration and classifiers

To develop the calibration and classifiers for UV-sensitive applications, the same analysis described in Sections 2.5 and 2.6 was conducted without AFMI channels with excitation at the UV range (excitation wavelengths from 400nm to 495nm).

## 2. RESULTS AND DISCUSSION

Linear regression combined with the method of swarm intelligence detected an approximately linear dependence between an optimized autofluorescence feature (spectral variable) and ROS levels determined by CellROX staining. This spectral variable was different for each sample and we did not find a significant similarity between spectral variables associated with different cell lines and tissue. The goodness of the calibration curves was statistically evaluated based on coefficient of determination (R) and associated p-values (Supplementary note 3 and supplementary table 2), which were found to range from 0.76 (p=0.00) to 0.92(p=0.00) for the full excitation spectrum, and 0.74 (p=0.00) to 0.87 (p=0.00) (Figure 2) for the UV-free spectrum (Supplementary Figure 1).

**Figure 2.**
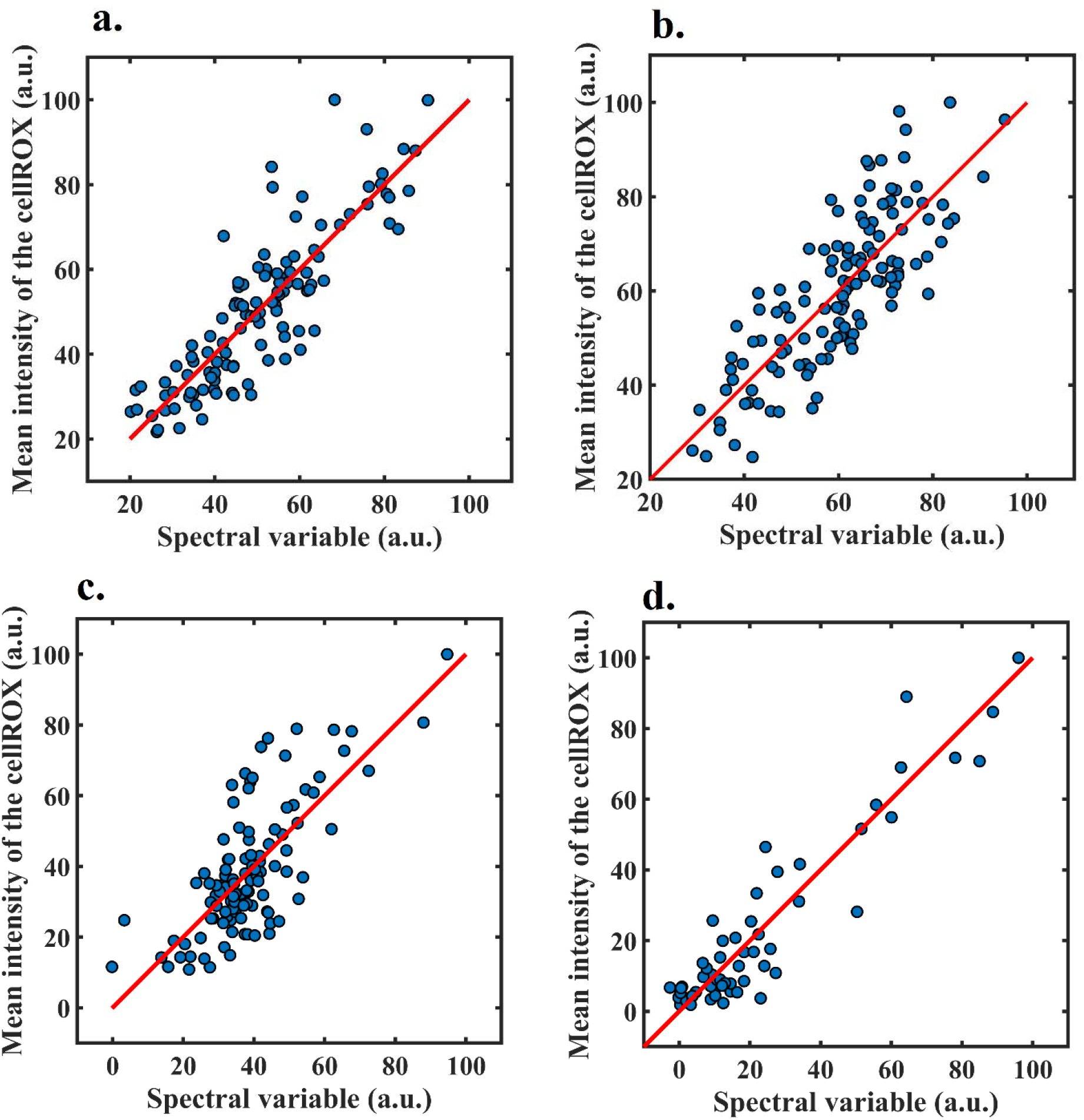
Regression curve used to calibrate the AFMI system to measure the states of the ROS using full spectrum. (a) HeLa (R=0.86), (b) PANC1 (R=0.83), (c) MSC (R=0.76) (d) kidney tissue (R=0.92)

We further developed classifiers to distinguish cells with a high level of ROS from those with a moderate level of ROS [29]. The same data and classification models developed here were used to evaluate the level of ROS in 3 different cell lines and kidney tissue. The classifiers were constructed using discrimination analysis to separate the cells with high-level ROS from cells with a moderate level [15](further details in supplementary note 4). Figure and Supplementary Figure 2 show the clusters formed by cells with high and moderate ROS level obtained using the entire excitation spectrum and UV-free excitation spectrum, respectively.

**Figure 3.**
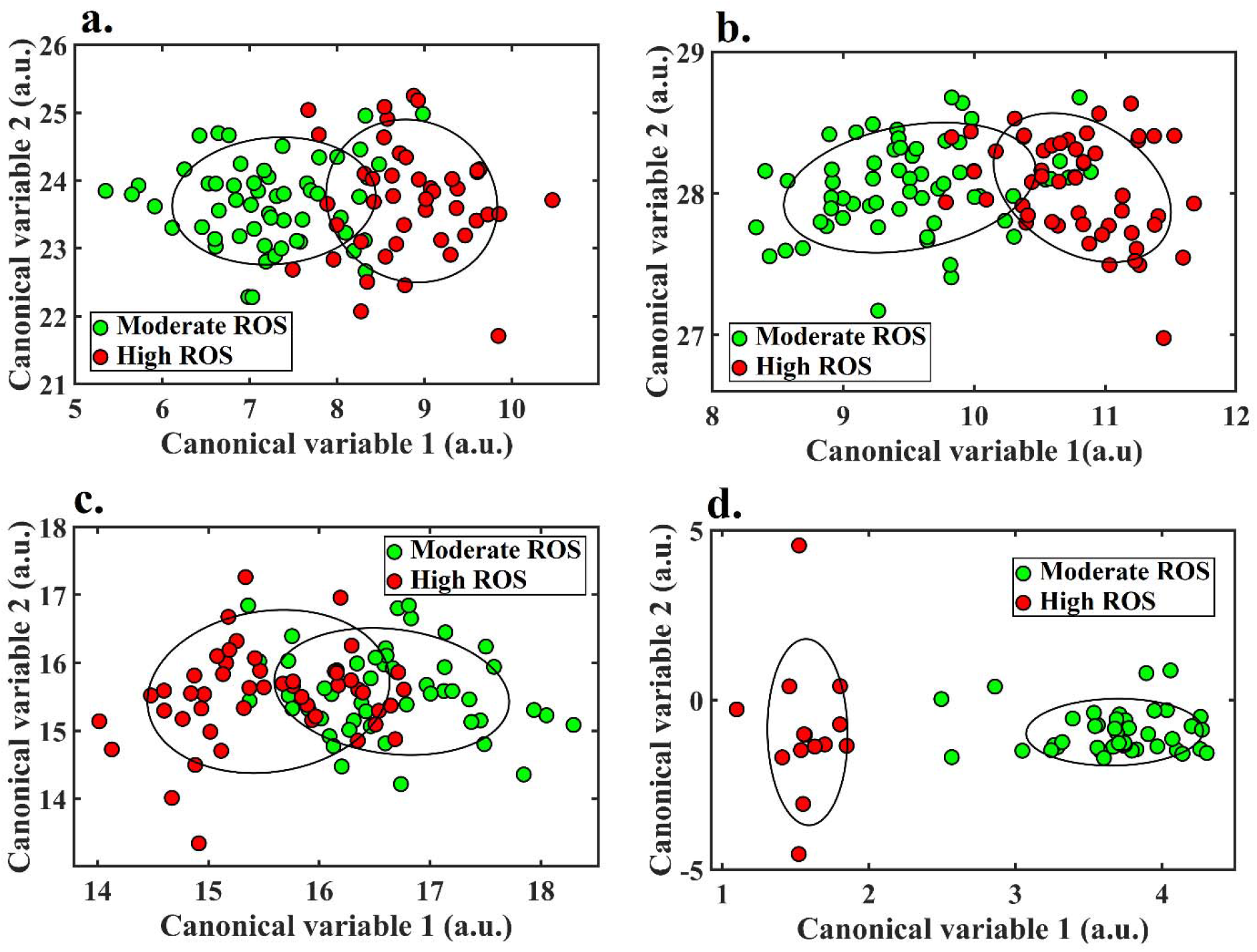
Cell/tissue classification into high and moderate ROS. To visualize the data distribution for each group, an ellipse was defined for each cluster which represents the standard deviation of the data points. (a) HeLa cells (IoU=8.3%), (b) PANC1 cells (IoU =5.1%), (c) MSCs (IoU=22%) (d) kidney tissue (IoU=0%). Results shown here used the full-spectral range.

The receiver operating characteristic (ROC) graph was obtained to determine the performance of this classifier for each sample as shown in Figure 4 [30]. Overall, the classifiers for different samples showed similar performance with average accuracies of 84% and 79% and AUCs of 0.91 and 0.87 for the entire excitation spectrum and UV-free excitation spectrum, respectively (see Supplementary Table 2 for further related statistics).

**Figure 4.**
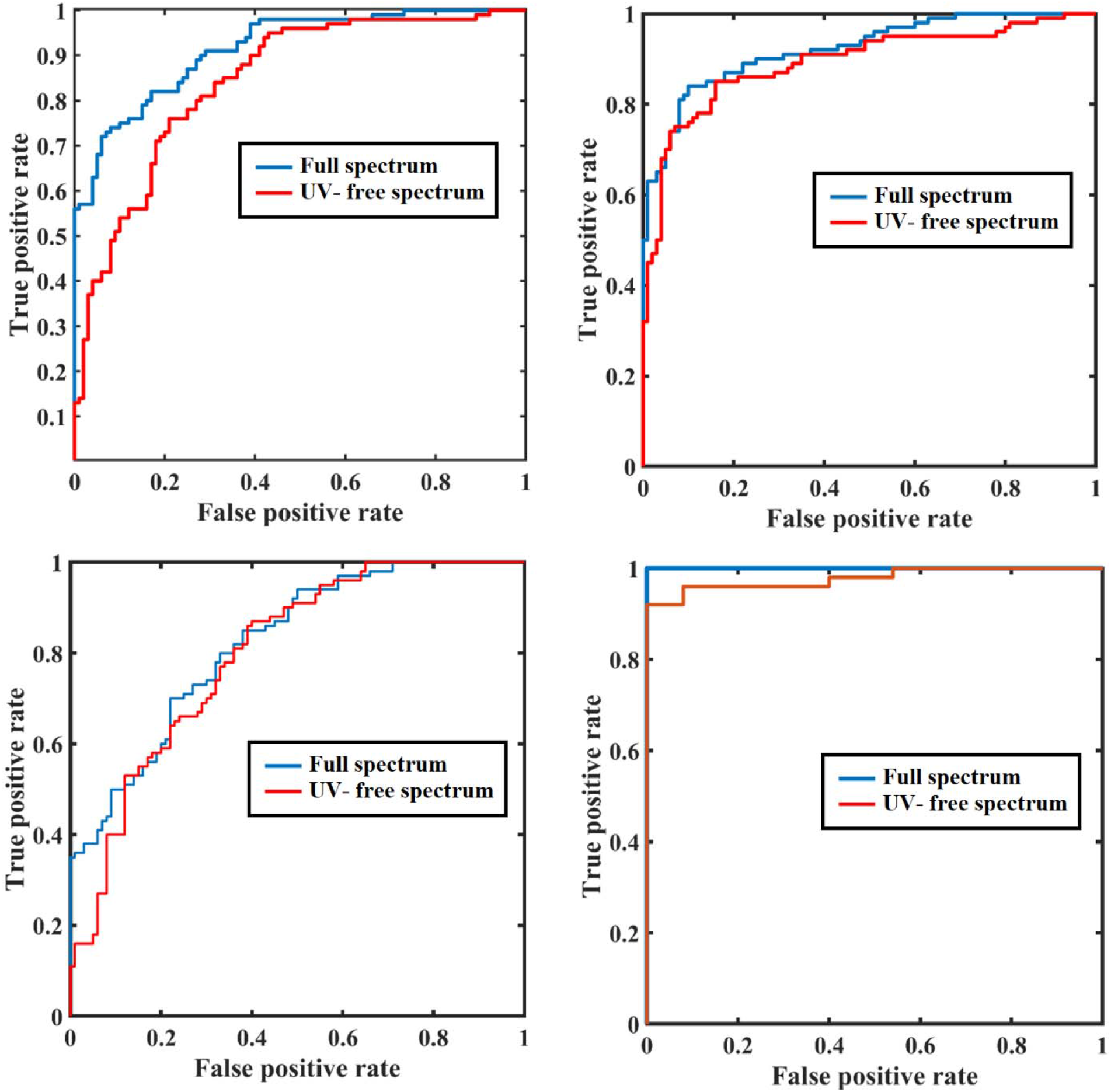

ROC curve for the accuracy of discrimination between cells with different level of ROS (a) HeLa cells (AUC=0.92/0.84 for full spectrum/UV-free spectrum), (b) PANC1 cells (AUC=0.92/0.89), (c) MSCs (AUC=0.82/0.80) (d) CKD tissue (AUC=1.00/0.98)

In this study, we demonstrate the use of a non-invasive auto-fluorescence multispectral imaging technique for the quantitative analysis of ROS-related spectral signatures of cells and tissue preserving spatial information. This technology is capable of measuring the value of ROS with high accuracy and also classifying the cells into moderate and high levels of ROS. It does not perturb biology as it does not require cell labeling or any other interference, and it has the ability to assess cryopreserved and fixed tissue, reflecting the redox state of cells after fixation.

Oxidative stress is a hallmark of multiple chronic diseases associated with older age such as diabetes, cardiovascular, renal, pulmonary, neurological and skeletal muscle disorders [31]. In addition, it is known to play a pivotal role in disease onset and development [32], such as the development and progression of diabetes complications both microvascular and macrovascular [33, 34]. As such, our novel technique has great clinical application not only in disease detection but also in studying the underlying mechanisms and role of oxidative stress in disease onset and progression.

AFMI requires a fairly simple adaptation of a fluorescence microscope using low power LEDs (see Supplementary Material Table 1) [15]; this allows the extraction of informative spectral maps of cells and tissues which contain details on the biochemical composition of the sample which correlates with the level of ROS. This simplicity and low cost make AFMI easily translatable to the clinical setting. The immediate application of this study would be monitoring ROS state in embryology, reproductive medicine and ophthalmology.

## 3. CONCLUSION

In conclusion, we found that cellular autofluorescence imaging with spectral selectivity non-invasively determines the level of ROS in cells and tissues. We developed calibration curves and classifiers for automatic measurement. AFMI provides a promising measurement tool for ROS which can be translated to future clinical and research applications.

## Acknowledgment

We acknowledge the support of the Australian Research Council CE140100003, and DP170101863, and National Health and Medical Research Council APP1144619. We thank Hui Chen (University of Technology Sydney) and Professor Stan Gronthos (South Australia Health and Medical Research Institute) for supplying the cells and animal tissue for the study.

