## supplementary material for "Non-invasive real-time imaging of reactive oxygen species (ROS) using multispectral auto-fluorescence imaging technique: a novel tool for redox biology"

**Suplimetary materials:**

### Supplementary note 1: Cell culture and tissue preparation

Cancer cell lines, HeLa and PANC1 (Sigma), were cultured in Dulbecco's modified Eagle's medium (DMEM) High glucose with L-glutamine (Gibco) with 10% fetal calf serum (FCS; Gibco) and Antibiotic-Antimycotic (Gibco). Human mesenchymal stem cells (MSCs), including ND0094 and ND0112, were isolated from bone marrow aspirates from the posterior iliac crest of informed donors aged 18-35 years following the provision of informed consent according to the approved procedures of the Human Ethics Committee of the Royal Adelaide Hospital, South Australia (ethics protocol number 940911a). In this study, data obtained from ND0094 and ND0112 were analysed together as mesenchymal stem cells (MSC). Multipotent STRO-1+ cells were isolated by immunomagentic selection and characterized as previously described [1, 2]. Their use in the current project was approved by the low-risk Human Ethics Committee of the University of New South Wales (ethics protocol number HC180219). MSCs were maintained in α-MEM with sodium bicarbonate, without L-glutamine, ribonucleosides and deoxyribonucleosides (Sigma) supplemented with 10% fetal bovine serum (Hyclone), sodium pyruvate (Sigma), L-Glutamine (Gibco), L-Ascorbic acid (Sigma, A8960) and Penicillin-Streptomycin (Sigma). All cultures were maintained in a humidified incubator at 5% CO2, 37$^{o}$C. The detachment of cells from culture surfaces was carried out with TrypLE (Gibco) after washing twice with Dulbecco's PBS without calcium or magnesium (Sigma). TrypLE was inactivated by adding twice its volume of growth media.

**Supplementary Table 1. Spectral channels and their specifications used in this study**

| Channel No. | Excitation wavelength ±5nm | Emission wavelength (bandwidth), nm | Dichroic miror Long pass, nm | Power at objective, uW |
|---|---|---|---|---|
| 1 | 334 | 447 (60) | 409 | 0.5 |
| 2 | 365 | 447 (60) | 409 | 1.1 |
| 3 | 375 | 447 (60) | 409 | 1.8 |
| 4 | 334 | 587 (35) | 532 | 0.1 |
| 5 | 365 | 587 (35) | 532 | 1.5 |
| 6 | 375 | 587 (35) | 532 | 11.5 |
| 7 | 385 | 587 (35) | 532 | 11.3 |
| 8 | 395 | 587 (35) | 532 | 19.4 |
| 9 | 405 | 587 (35) | 532 | 23.5 |
| 10 | 415 | 587 (35) | 532 | 34.0 |
| 11 | 425 | 587 (35) | 532 | 62.6 |
| 12 | 435 | 587 (35) | 532 | 85.9 |
| 13 | 455 | 587 (35) | 532 | 40.3 |
| 14 | 475 | 587 (35) | 532 | 102.7 |
| 15 | 495 | 587 (35) | 532 | 43.1 |
| 16 | 405 | 700 (long pass) | 635 | 23.9 |
| 17 | 455 | 700 (long pass) | 635 | 41.4 |
| 18 | 495 | 700 (long pass) | 635 | 94.8 |

### Supplementary note 2: Spectral variable discovery for the calibration curve

The spectral variable was optimally discovered in this study. This is traditionally achieved by using linear regression, impossible in this case due to a large number of features (324) leading to overfitting. We, therefore, used the method of “swarm intelligence” [3] which allows the identification of optimal smaller feature sets (no more than 15 features in this case) for linear regression, thus avoiding overfitting. The fundamental building block in this method is linear regression between selected 15 features from the AFMI vector for each cell and the corresponding CellROX ROS value from that cell. The application of linear regression returns a line of best fit, in a relevant 16-dimensional space where the selected 15 features serve as independent variables and the ROS feature acts as the dependent variable. We also obtain the value of the error term $\varepsilon_I$ for these 15 features, in this case the sum of squared errors for each tested cell in the linear regression.

The method of swarm intelligence was applied here to identify the optimized 15 features. In this method, multiple agents independently move about in an abstract space specific to the problem under consideration. As they move, they attempt to optimize a set criterion whose value depends on the location in that space. As we are attempting to optimize the 15 features simultaneously, the space in our case is the space S of all possible sets of 15 features. The space S is a discrete grid made of individual points $N_I = (n_1, n_2, \dots n_{15})$, where $n_i$ $(i = 1, \dots 15)$ is a feature number, from 1 to 342. As we have 342 available features, the number of points in this grid, equal to the number of possible tens of features is $342^{15}$. As indicated previously, each grid point $N_I$ is characterized by the error term $\varepsilon_I$.

In the first step of our method, we randomly chose 50 grid points, $N_{ini}$ from the space S as the locations of our agents forming the swarm; these agents move synchronously, each on their own trajectory, according to the rule we have imposed on their movement in the space S. This rule first identifies the agent and its grid point $N_{min}$ where the error term $\varepsilon_{min}$ is the smallest overall (in the entire space S). For each agent, the rule also identifies the minimal error term over this agent's trajectory $\varepsilon_{loc}$ and the associated grid point on this trajectory, $N_{loc}$. These three grid points, $N_{ini}$, $N_{loc}$ and $N_{min}$ determine where the agent will go next. This location is the nearest grid point to the vector $\alpha_1 N_{ini} + \alpha_2 N_{loc} + \alpha_3 N_{min}$ where we have chosen $\alpha_{1,2,3}$ to be 0.15.

The agents are allowed to move until the agent's trajectories converge at a final grid point, which is the location of the optimized fifteen features. We note that there is no mathematical guarantee that the minimum obtained through swarm intelligence is a global minimum – as we are only sampling a part of the space. However, the sampling is comprehensive and reasonably random, and this increases the chances that the detected minimum is indeed global [3]. The results of linear regression using these optimized 15 features is shown in Figure 1.

### Supplementary note 3: Calibration curve statistical tests

To statistically evaluate the goodness of fit associated with the calibration curve (Figure 2), the coefficient of determination (R), and P-values (Supplementary **Error! Reference source not found.**) were obtained. The coefficient of determination (R) evaluates how closely the predicted ROS value from the regression model matches the reference ROS values which varies between 0 to 1 from the least to the best fitting. Overall, the R-value for full spectrum excitation and UV-sensitive excitations are 0.84 and 0.78, respectively. "P-value" evaluates the overall significance of the regression model which assesses the null hypothesis that all of the regression coefficients are equal to zero. A P-value less than the default significance level of 0.05 is the indication of a significant linear regression relationship between the ROS value and spectral variable. Supplementary **Error! Reference source not found.** shows that the p-value for all regression curves is 0.00 which demonstrates the significance of all curves for both full excitation spectrum and UV sensitive spectrum.

**Supplementary Table 2. Statistical tests of regression curves**

| Cell line | Excitation range | R | P-value |
|---|---|---|---|
| Hela | 350-600 | 0.86 | 0.00 |
| | 400-600 | 0.74 | 0.00 |
| PANC1 | 350-600 | 0.83 | 0.00 |
| | 400-600 | 0.78 | 0.00 |
| MSC | 350-600 | 0.76 | 0.00 |
| | 400-600 | 0.75 | 0.00 |
| Kidney tissue | 350-600 | 0.92 | 0.00 |
| | 400-600 | 0.87 | 0.00 |

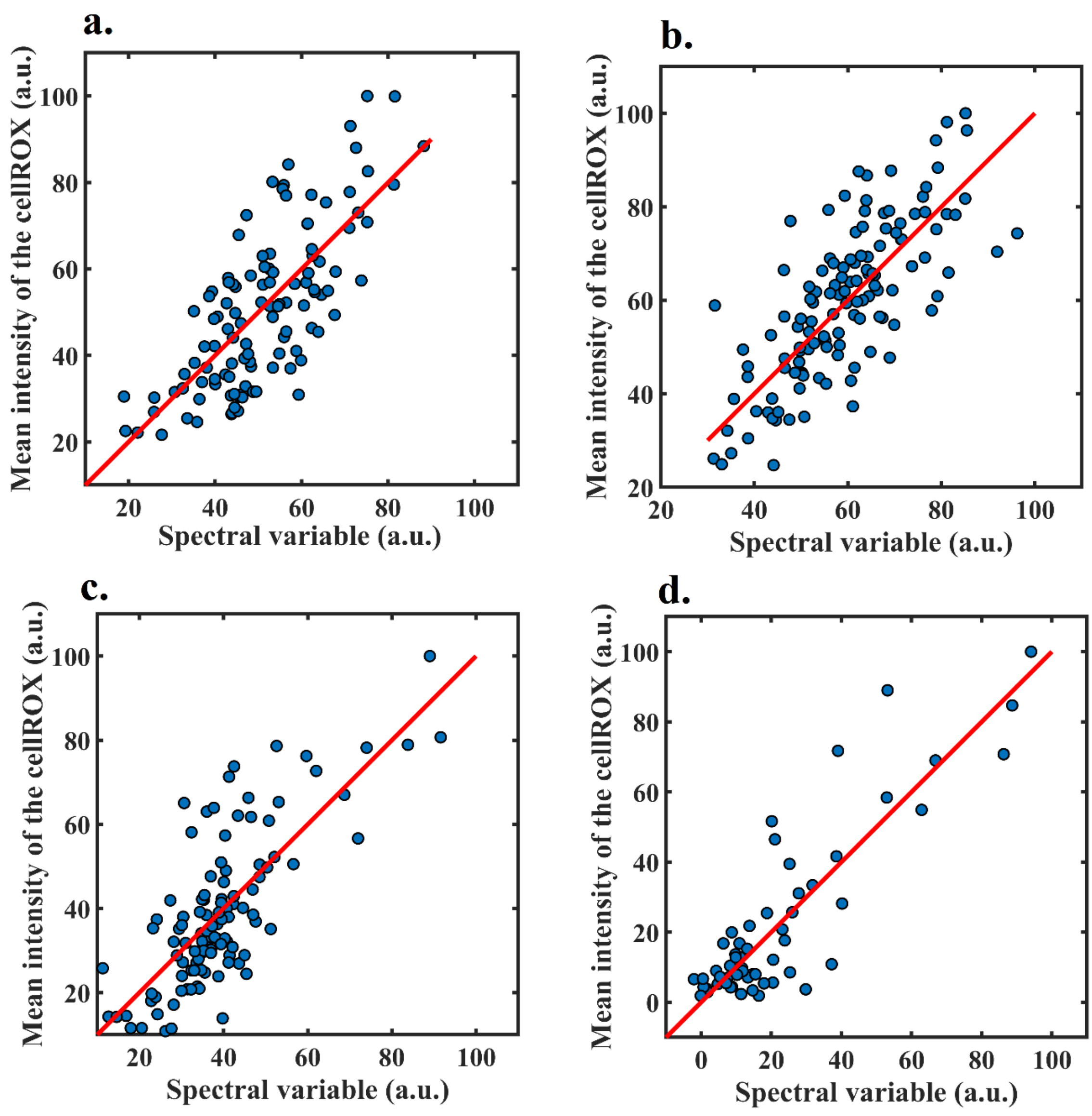


Supplementary Figure 1. Regression curve used to calibrate the AFMI system to measure the states of the ROS using UV-free spectrum. (a) HeLa (R=0.74), (b) PANC1 (R=0.78), (c) MSC(R=0.75) (d) CKD(R=0.87)

### Supplementary note 4: Classification and cluster discovery

Discriminative analysis was employed to obtain the optimal separation between high and moderate ROS groups. The color features from cell images including mean cell image intensity and their ratio were obtained. The correlated features were identified using the Pearson correlation test and removed. Further, the 15 most indicative features ($P < 0.001$) were selected as an AFMI feature vector to feed to the discrimination analysis.

The optimal separation of the high and moderate ROS groups was obtained by reducing the AFMI feature vectors dimension  projecting the onto an optimal two-dimensional (2-D) space created by discriminative analysis [4]. This space maximizes between-group distance while minimizing within-group variance. The projection reduces the dimension of the AFMI vectors to two, and it determines canonical variables which are equal to the linear combination cellular features obtained by this projection [5]. These spaces were different for each sample compared due to utilizing different features vectors, as different features were indicative for different samples.

To quantify the overlap of clusters quantitatively, intersection over union (IoU) values were calculated [6]. IoU is the ratio of the area of the two-ellipse intersections, divided by the area of their union which varies from 0% to 100% for fully separated to fully overlapped ellipses, respectively. IoU analyses demonstrated that samples with different levels of ROS form separate clusters while a degree of overlap persists (Supplementary Table 2). The average IoU value for separating cells with high level of ROS from moderate level of ROS was found to be 8.85% and 14.8% for full-spectrum and UV-sensitive spectrum, respectively.

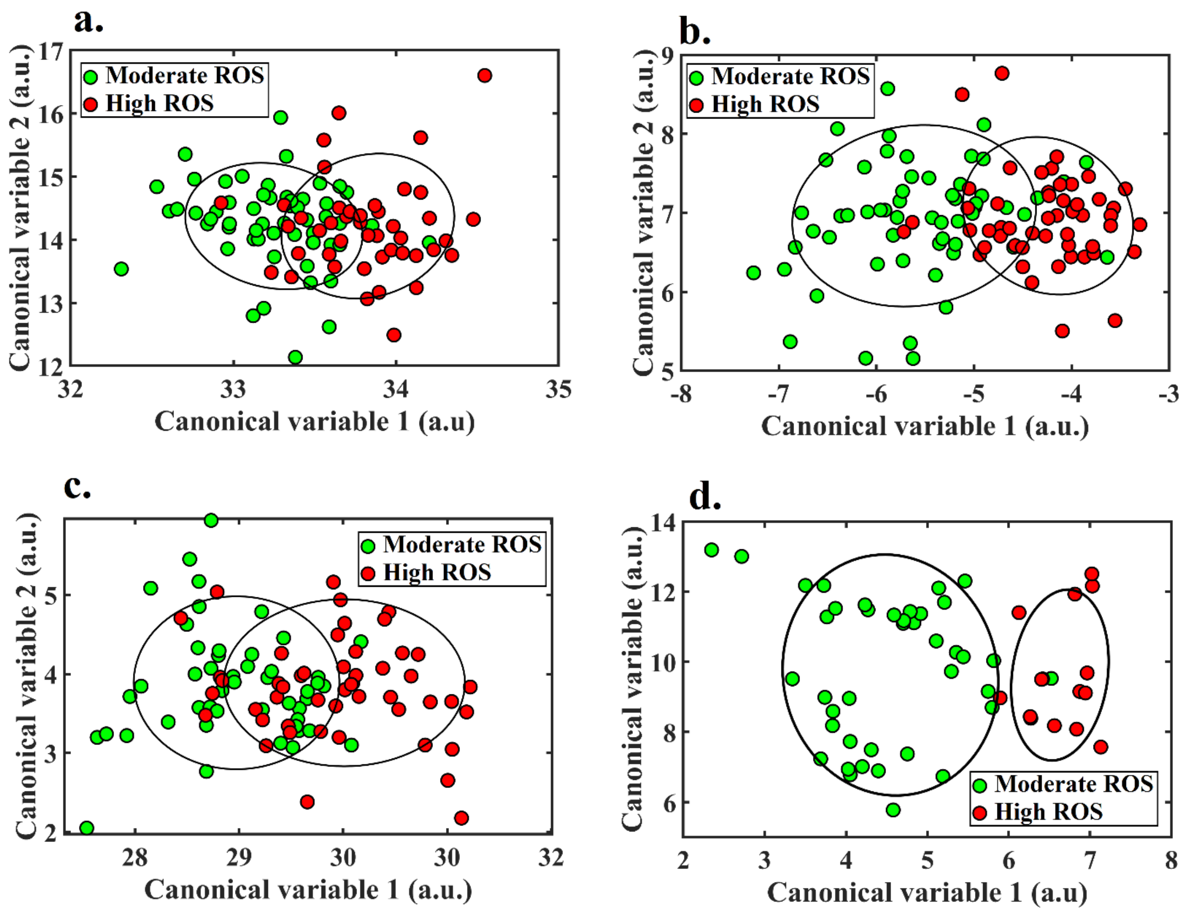

Supplementary Figure 2. Sample classification into high to moderate ROS. To visualize the data distribution for each group, an ellipse was defined for each cluster which represents the standard deviation of the data points. (a) HeLa (IoU=20%), (b) PANC1 (IoU =13%), (c) MSC(IoU=26%) (d) CKD ( IoU=0%). Results shown are for the UV-free spectral range

Further, a linear classifier was employed to predict the pre-defined cell level of ROS [7, 8]. To evaluate the classifier, we used a cross-validation methodology [9] wherein data points were partitioned into 10 groups, a linear classifier was developed based on 9 of these groups, and the tenth group was used for testing and the calculation of accuracy. The receiver operating characteristic (ROC) graph was obtained to determine the performance of this classifier as its discrimination threshold varied [10, 11].

Supplementary Table 3. Classification performance measures

| Sample | Excitation range | IoU | AUC | Accuracy |
|---|---|---|---|---|
| Hella | 350-600 | 8.3% | 0.92 | 82% |
| | 400-600 | 20% | 0.84 | 77% |
| PANK | 350-600 | 5.1% | 0.92 | 86% |
| | 400-600 | 13.2% | 0.89 | 79% |
| MSC | 350-600 | 22% | 0.82 | 72% |
| | 400-600 | 26% | 0.80 | 68% |
| Kidney tissue | 350-600 | 0% | 1.00 | 98% |
| | 400-600 | 0% | 0.98 | 93% |
